# Mutational landscape of a chemically-induced mouse model of liver cancer

**DOI:** 10.1101/242891

**Authors:** Frances Connor, Tim F. Rayner, Christine Feig, Sarah J. Aitken, Margus Lukk, Duncan T. Odom

**Affiliations:** Cancer Research UK Cambridge Institute, University of Cambridge, Robinson Way, Cambridge CB2 0RE, UK.; Department of Histopathology, Addenbrooke’s Hospital, Cambridge University Hospitals NHS Foundation Trust, Hills Road, Cambridge CB2 0QQ, UK.

**Keywords:** Hepatocellular carcinoma, carcinogen, exome sequencing, mutational signatures, *Hras*

## Abstract

Carcinogen-induced mouse models of liver cancer are used extensively to study the pathogenesis of the disease and have a critical role in validating candidate therapeutics. These models can recapitulate molecular and histological features of human disease. However, it is not known if the spectra of genomic alterations driving these mouse tumour genomes are comparable to those found in humans. Here, we provide a detailed characterisation of the exome-wide pattern of mutations in tumours from mice exposed to diethylnitrosamine (DEN), a commonly used model of hepatocellular carcinoma (HCC). DEN-initiated tumours had a high, uniform number of somatic single nucleotide variants (SNVs), with very few insertions, deletions or copy number alterations, consistent with the known genotoxic action of DEN. Exposure of hepatocytes to DEN left a reproducible mutational imprint in resulting tumour exomes which we could computationally reconstruct using six known COSMIC mutational signatures. The tumours carried a high diversity of low-incidence, non-synonymous point mutations in many oncogenes and tumour suppressors, reflecting the stochastic introduction of SNVs into the hepatocyte genome by the carcinogen. We identified four recurrently mutated genes that were putative oncogenic drivers of HCC in this model. Every neoplasm carried activating hotspot mutations either in codon 61 of *Hras*, in codon 584 of *Braf* or in codon 254 of *Egfr*. Truncating mutations of *Apc* occurred in 21% of neoplasms, which were exclusively carcinomas supporting a role for deregulation of Wnt/β-catenin signalling in cancer progression. ***Conclusion:*** Our study provides detailed insight into the mutational landscape of tumours arising in a commonly-used carcinogen model of hepatocellular carcinoma, facilitating the future use of this model to understand the human disease.

## Introduction

Hepatocellular carcinoma, the predominant form of primary liver cancer, is currently the sixth most frequently diagnosed human cancer. HCC is the second most common cause of cancer death globally and its incidence is increasing in countries with historically low rates (1, 2). HCC typically develops in the context of end-stage liver disease, resulting from chronic inflammation, fibrosis and cirrhosis, and is almost exclusively caused by environmental risk factors, such as chronic hepatitis virus infection, aflatoxin B exposure, chronic alcohol consumption, and metabolic syndrome (3). This diversity of aetiologies appears to be reflected in the molecular heterogeneity of the disease. Over the last few years, next generation sequencing analyses of hundreds of human liver tumours have identified several oncogenic pathways and a wide range of putative driver gene mutations underlying hepatocarcinogenesis (4–9).

There are an increasing number of experimental mouse models used in HCC research to study the disease pathogenesis and to assess novel therapeutics (10). For several decades, carcinogen-induced tumours have been used in preclinical research, and the most widely used chemical to induce liver cancer in mice is diethylnitrosamine. When injected into juvenile mice, DEN targets the liver where it is metabolically activated by centrilobular hepatocytes into alkylating agents that can form mutagenic DNA adducts (11). The introduction of oncogenic mutations into hepatocytes that are actively proliferating during normal post-natal development can then result in dysplastic lesions which progress to carcinoma. Mouse tumours induced by DEN alone frequently harbour initiating activating mutations in either *Hras* or *Braf* proto-oncogenes (12, 13). In a related model in which tumours are induced using DEN as an initiator followed by phenobarbital as a tumour promoter, chromosomal instability and activating mutations in β-catenin have been implicated in tumour progression (14). There is also evidence that inflammation is a contributing factor to DEN-induced hepatocarcinogenesis. As well as acting as a genotoxin, DEN is also hepatotoxic causing necrotic cell death. This damage triggers an inflammatory response resulting in elevated expression of mitogens, such as interleukin-6, which promote compensatory proliferation of surviving hepatocytes (15).

No single mouse model can capture all aspects of human HCC, although each can recapitulate at least some of the genetic and/or cellular features of the human disease. For example, a comparison of global gene expression profiles showed that HCC from DEN-treated mice resembles a subclass of human HCC associated with poor prognosis (16). However, there are few studies which compare the genome-wide mutational landscapes of mouse cancer models to those seen in the human cancer. Such oncogenomic evaluations will be crucial for identifying the most appropriate preclinical mouse model for specific clinical questions. To this end, we describe the exome-wide mutational pattern in tumours arising in the DEN mouse model of HCC, which has, and continues to be, commonly used in preclinical research to understand the biology of liver cancer.

## Materials and Methods

Extended materials and methods can be found in the Supplementary Information.

## Generation of mouse samples

Male C3H/HeOuJ male mice were administered a single intraperitoneal injection of DEN (20mg/kg body weight) aged 14 to 16 days. Liver samples were collected during the first 24 hours following DEN administration; tumour samples were collected up to 40 weeks after treatment. Untreated C3H male mice were used for reference tissue samples or aged up to 76 weeks for spontaneous liver tumour samples. Tissue samples were snap frozen for DNA extraction and/or fixed in neutral buffered formalin for histological analyses.

## Histological analyses

Histochemical staining with haematoxylin and eosin (H&E) or using the Gomori’s method was carried out on formalin-fixed paraffin-embedded tissue sections. Immunohistochemistry was performed using antibodies against β-catenin (BD Biosciences); phospho-histone H2AX (Merck Millipore); O^6^-ethyl-2-deoxyguanosine (ER6, Squarix Biotechnology); and Ki67 (Bethyl Laboratories). Quantification of nuclear staining for O^6^-ethyl-2-deoxyguanosine and phospho-histone H2AX was done using ImageScope software (Leica Biosystems). Tumours were classified according to the International Harmonization of Nomenclature and Diagnostic Criteria for Lesions in Rats and Mice (INHAND) guidelines (17).

## DNA isolation, whole exome sequencing and sequence alignment

Genomic DNA was isolated from liver tumours and from ear/tail samples using the AllPrep DNA/RNA mini kit or the DNeasy blood & tissue kit (Qiagen), according to the manufacturer’s instructions. Exome capture libraries were prepared following the instructions of the SureSelectXT mouse all exon target enrichment system (Agilent Technologies). Exome libraries were sequenced using a 125bp paired-end read protocol on an Illumina HiSeq 2500.

Sequencing reads were aligned to the C3H_HeJ_v1 mouse genome assembly (Ensembl release 90 (18)) using BWA (versions 0.6.1 or 0.7.12 (19)). Aligned bam files were annotated using Picard tools (version 1.124; http://broadinstitute.github.io/picard) and sequencing coverage metrics calculated using samtools (version 1.1 (20)). Aligned reads for human samples from the LICA-FR and LIAD-FR cohorts were downloaded from the European Genome-phenome Archive at EMBL-EBI (accessions EGAD00001000131, EGAD00001001096 and EGAD00001000737).

## Variant identification, prioritisation and validation

Single nucleotide and indel variants were called using Strelka (version 1.0.14 (21)) and autosomal copy number variants were called using CNVkit (version 0.7.2 (22)). Variants were subject to multiple filtering steps, as described in the Supplementary Information.

High-likelihood cancer driver genes were prioritised from a list of genes known to harbour *bona fide* cancer driver mutations (23). Cancer genes with above expected levels of non-synonymous mutations were identified by fitting the number of observed mutations at each variant locus to a Poisson distribution, and then combining variant loci at the gene level using a multinominal model from the R XNomial package (version 1.0.4; https://CRAN.R-project.org/package=XNomial). Non-synonymous SNVs in the identified cancer driver genes of interest, *Hras, Braf, Egfr* and *Apc*, were confirmed using conventional Sanger sequencing or by visual inspection of aligned reads.

## Phylogenetic and mutational signature analyses

A phylogenetic tree was built in R using the ape package (version 3.5 (24)). Pairwise distances between samples were calculated as the number of genomic loci at which the sample genotypes differ and trees were constructed using a neighbour-joining algorithm (25).

For the mutational spectra analysis, SNVs were annotated by the 96 possible trinucleotide context substitutions. The distributions of 5' and 3' nucleotides flanking the SNVs were calculated directly from the reference genome. Comparison between human and mouse mutational signatures was facilitated by normalisation of C3H/HeJ nucleotide context distributions using the ratios of known trinucleotide prevalences in C3H/HeJ and human genomes. The proportions of COSMIC mutational signatures (26) represented in the mutational profile from each sample was calculated using the R package deconstructSigs (version 1.8.0 (27)).

## Results

### DEN-initiated carcinogenesis in mouse hepatocytes

We generated chemically-initiated liver tumours using a well-established protocol in which juvenile (14–16 days old) C3H/HeOuJ male mice were administered a single intraperitoneal injection (20mg/kg body weight) of diethylnitrosamine (28). Animals were then aged for up to 40 weeks. The C3H strain was chosen because it is highly susceptible to the development of both treatment-induced and spontaneous liver tumours (29). We therefore also aged a cohort of untreated C3H male mice for up to 76 weeks to generate spontaneous liver neoplasms for comparison with the DEN-initiated tumours (Fig. 1A).

**Figure 1.**
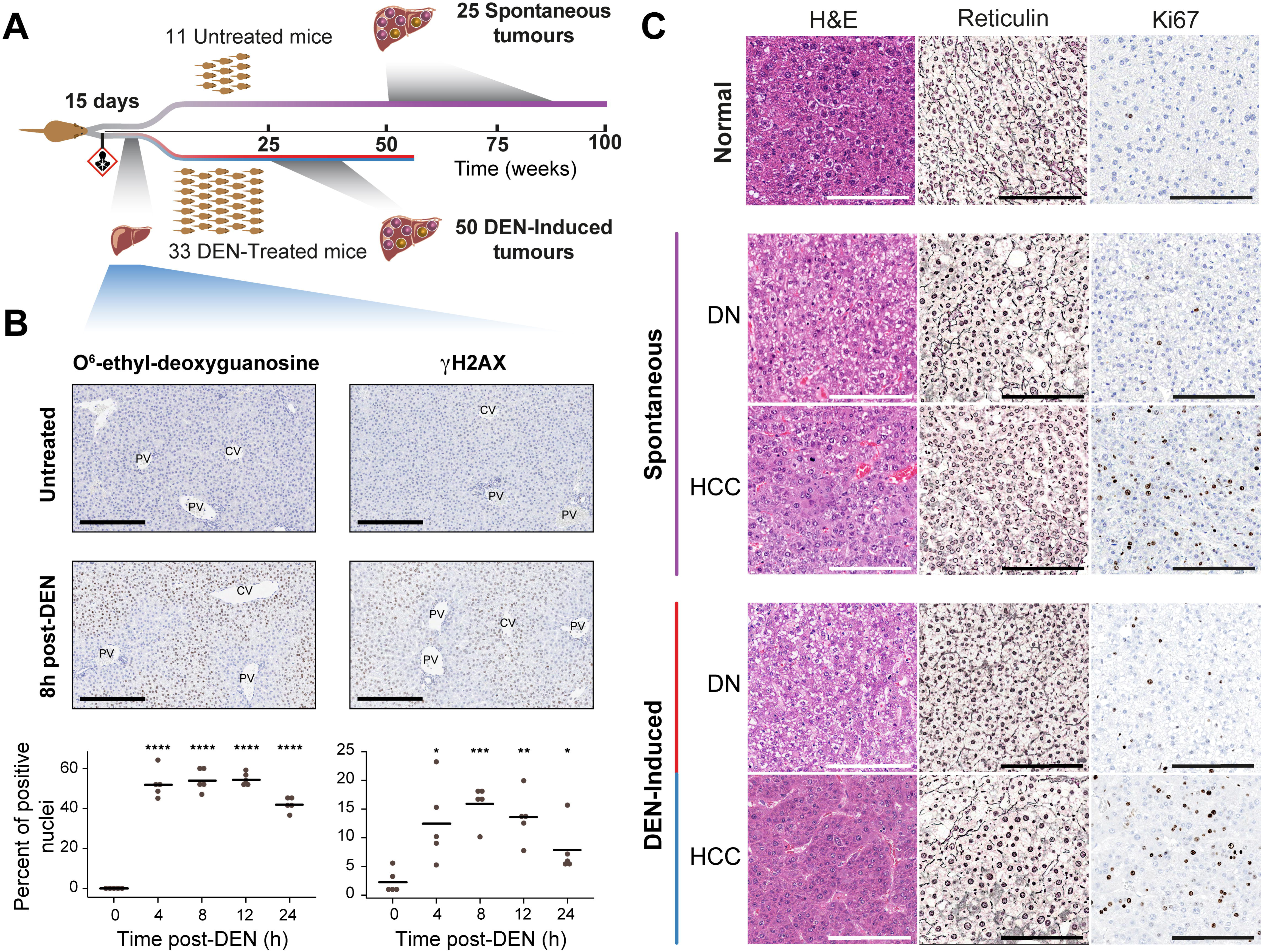
DEN-initiated carcinogenesis in mouse hepatocytes. (A) Overview of experimental design. Cohorts of C3H male mice were either administered DEN aged 14–16 days or left untreated. Liver samples were collected during the first 24 hours after DEN exposure for histopathological analysis, or mice were aged to develop tumours. Dysplastic nodules and HCCs were collected from cohorts of mice 24–26 weeks and 26–40 weeks, respectively, after administration of DEN; spontaneous tumours were collected from mice aged 37–76 weeks. (B) *In situ* detection and quantification of DEN-induced DNA damage. Representative photomicrographs of immunohistochemistry for O^6^-ethyl-2-deoxyguanosine and gamma-H2AX in untreated mice and 8 hours post-DEN injection (PV: portal vein; CV: central vein). All scale bars = 200[jm. Original magnification x200. Automated image quantification was used to evaluate the dynamics of DNA damage for O^6^-ethyl-2-deoxyguanosine and gamma-H2AX over the course of 24 hours (n=5 samples per timepoint; bar indicates mean; Welch two-sample t-test: * p < 0.05, ** p < 0.01, *** p<0.001, **** p<0.0001). (C) Histology of murine hepatocellular neoplasms. Representative photomicrographs of serial sections of normal liver tissue and liver tumours arising in DEN-treated and untreated mice (DN: dysplastic nodule; HCC: hepatocellular carcinoma). H&E staining demonstrates tissue morphology; reticulin staining is used to assess architecture (normal: staining around each cord of hepatocytes; DN: loss of regular architecture; HCC: thickened trabeculae and corresponding reduction in staining); and Ki67 identifies mitotic cells (normal adult liver is mostly quiescent; DN and HCC show increasing numbers of dividing cells). All scale bars = 200μm. Original magnification x200.

DEN primarily targets the liver, in which it is metabolically activated by cytochrome P450 enzymes in hepatocytes (30). The resulting DEN metabolites can directly damage DNA by alkylating nucleobases. Of particular interest are the O6 position in guanine and the O4 position in thymine, which are both vulnerable to nucleophilic attack resulting in adducts with the potential to be miscoding (11). We examined the immediate DNA damage in 15 day old C3H livers over 24 hours after exposure to DEN. Metabolic activation of DEN occurred within 4 hours after administration, as seen by the presence of the promutagenic O6-ethyl deoxyguanosine adduct using immunohistochemistry (Fig. 1B). As expected, the majority of positively staining cells were found in centrilobular (zone 3) hepatocytes, consistent with the known high expression of cytochrome P450 enzymes and more extensive drug metabolism by hepatocytes in this region (31). As may be expected, DNA double strand breaks also accumulated after DEN treatment, as seen by the rapid accumulation and elimination of phosphorylated histone H2AX over the following 24 hours (Fig. 1B).

All DEN-treated mice developed multiple, macroscopically identifiable tumours by 25 weeks after administration, concordant with previous studies (32). H&E and reticulin stained tumour tissue sections from DEN-treated and untreated mice (Fig. 1C) were classified by a histopathologist using standardised INHAND diagnostic criteria (17); this revealed that all neoplasms had a hepatocellular phenotype. Almost all tumours arising in mice up to 26 weeks after DEN treatment were dysplastic nodules (DNs). Hepatocellular carcinomas were present at later time points, some of which had a nodule-in-nodule appearance, supporting the hypothesis of stepwise progression from DN to HCC (33). In addition to the macroscopically dissected tumours, examination of residual liver tissue of DEN-treated mice revealed microscopic basophilic and eosinophilic foci of cellular alteration (data not shown). These localised proliferations of phenotypically distinct hepatocytes represent potential neoplastic precursors to DNs, and in turn HCC (17).

The development of spontaneous tumours in untreated C3H showed greater histological and temporal variability (37–76 weeks). Importantly, DN and HCC tumours arising in these untreated mice were histologically indistinguishable from those treated with DEN (Fig. 1C). Furthermore, all these murine tumours histopathologically mimic their corresponding human tumours.

### Diversity of somatic SNVs reveals independent evolution of DEN-induced neoplasms

Whole exome sequencing was performed on DNA isolated from 50 discrete neoplasms excised from the livers of 33 individual C3H male mice given a single intraperitoneal administration of DEN as juveniles. 34 of the DEN-induced neoplasms were of sufficient size to provide additional tissue for histopathological examination; of these, 16 were classified as DNs and 18 as HCCs. The whole exome sequences of the remaining 16 DEN-induced neoplasms were used only for the phylogenetic analysis (see below and Fig. 2). In addition, whole exome sequencing was performed on DNA isolated from 25 macroscopically visible liver neoplasms (22 DNs and 3 HCCs) found in 11 untreated C3H male mice. The targeted exonic regions were sequenced to an average depth of 380x, with 95% of coding DNA sequences covered at >20-fold. Sequencing data were processed to identify somatic nucleotide substitutions, small insertion and deletion mutations and copy number alterations larger than 10MB.

**Figure 2.**
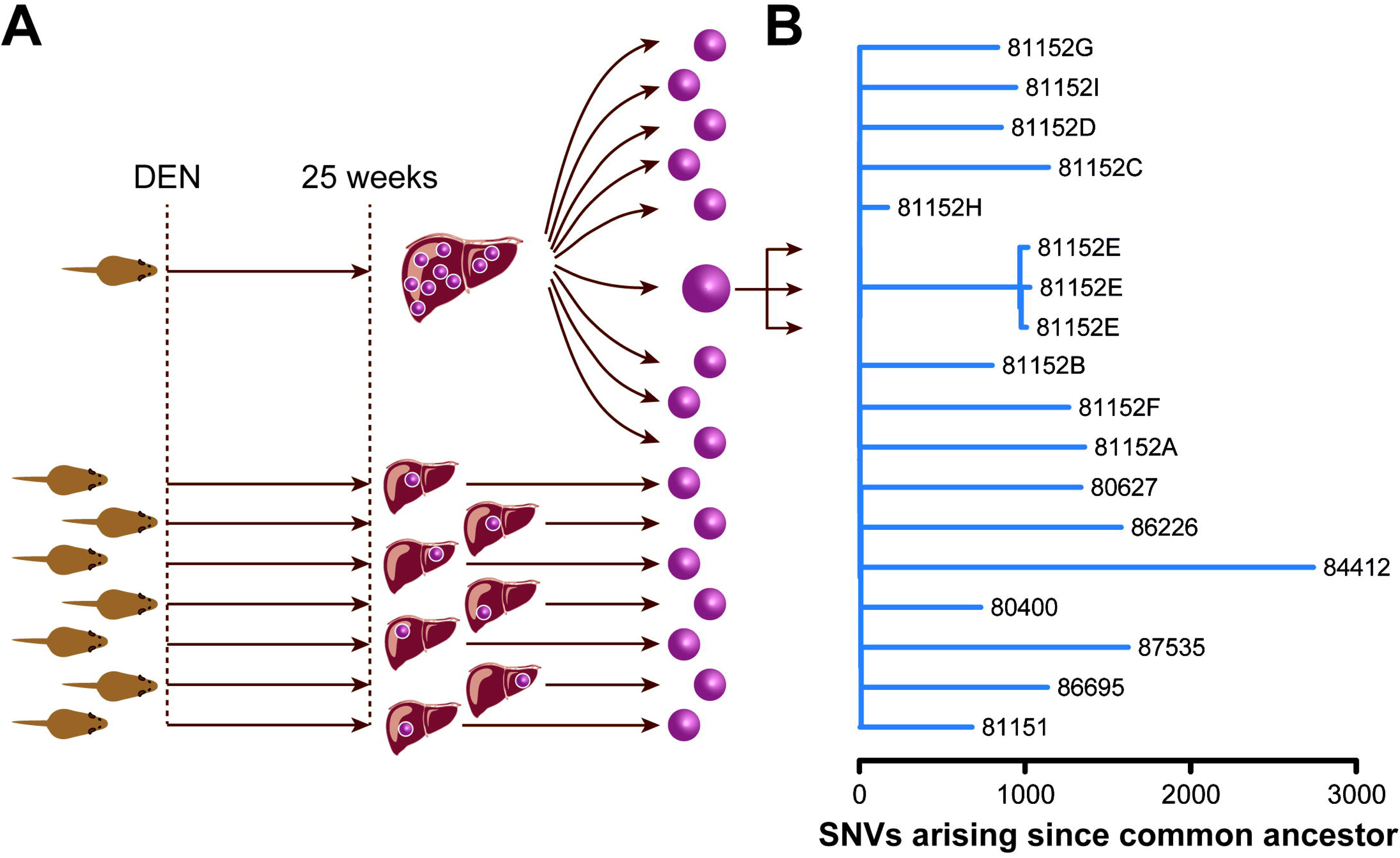
Independent evolution of DEN-initiated liver tumours is revealed by their unique SNV profiles. (A) Experimental design. Liver neoplasms were generated by intraperitoneal injection of DEN into 14–16 day old mice, which were then aged for 24–26 weeks. We performed whole exome sequencing of nine separate nodules isolated from a single mouse liver and of single nodules from livers of seven other mice. To evaluate the noise associated with library preparation, triplicate sequencing libraries were prepared in three separate batches for a single nodule. (B) Phylogenetic analysis of DEN-initiated tumours. A phylogenetic tree was constructed using the ape package in R, where branch lengths correspond to the number of unshared SNVs. Long branch lengths indicate no relatedness among the nodules within a single mouse, whereas three replicate libraries from the single tumour had short branches, indicating few SNV differences. Branches are labelled using mouse and tumour identification codes.

To test whether multiple tumours within one individual mouse treated with DEN had evolved independently, we constructed a phylogenetic tree to examine how closely related the mutational patterns were among nine nodules isolated from a single liver (Fig. 2). The DNA from one of these nodules was isolated and three separate libraries were generated to perform independent exome sequencing. In addition, seven nodules from seven different animals were also included where DEN-induced mutational patterns must have arisen autonomously. As expected, the three exome SNV profiles generated from the single nodule were almost identical. In contrast, very few of the 24,721 SNVs that we identified across all 16 samples in this cohort were shared between neoplasms. Indeed, the SNV profiles of separate neoplasms isolated from the same liver were as divergent as those isolated from separate mice, suggesting that each DEN-induced neoplasm was initiated, and evolved, as an independent tumour.

### Carcinogen-initiated liver tumours have a high SNV burden

Carcinogen-initiated neoplasms had reproducibly high numbers of somatic SNVs, with an average of 28.4 coverage-independent SNVs per Mb in histologically classified HCCs (Fig. 3A). Dysplastic nodules harboured fewer SNVs on average, albeit still at comparably high numbers (mean 22.1 per Mb). Despite sharing similar histology, the neoplasms which arose spontaneously in untreated mice had much lower SNV burdens, on average nineteen-fold fewer SNVs per Mb compared with the carcinogen-induced neoplasms. The lower numbers of SNVs in spontaneous tumours is comparable with those seen in human HCC (6). By contrast, both murine DEN-induced and spontaneous tumours carried very few somatic indels and copy number variants (Fig. 3B & C). The widespread acquisition of SNVs specifically in the exomes of DEN-induced neoplasms reflects the involvement of a DNA damaging chemical in their pathogenesis.

**Figure 3.**
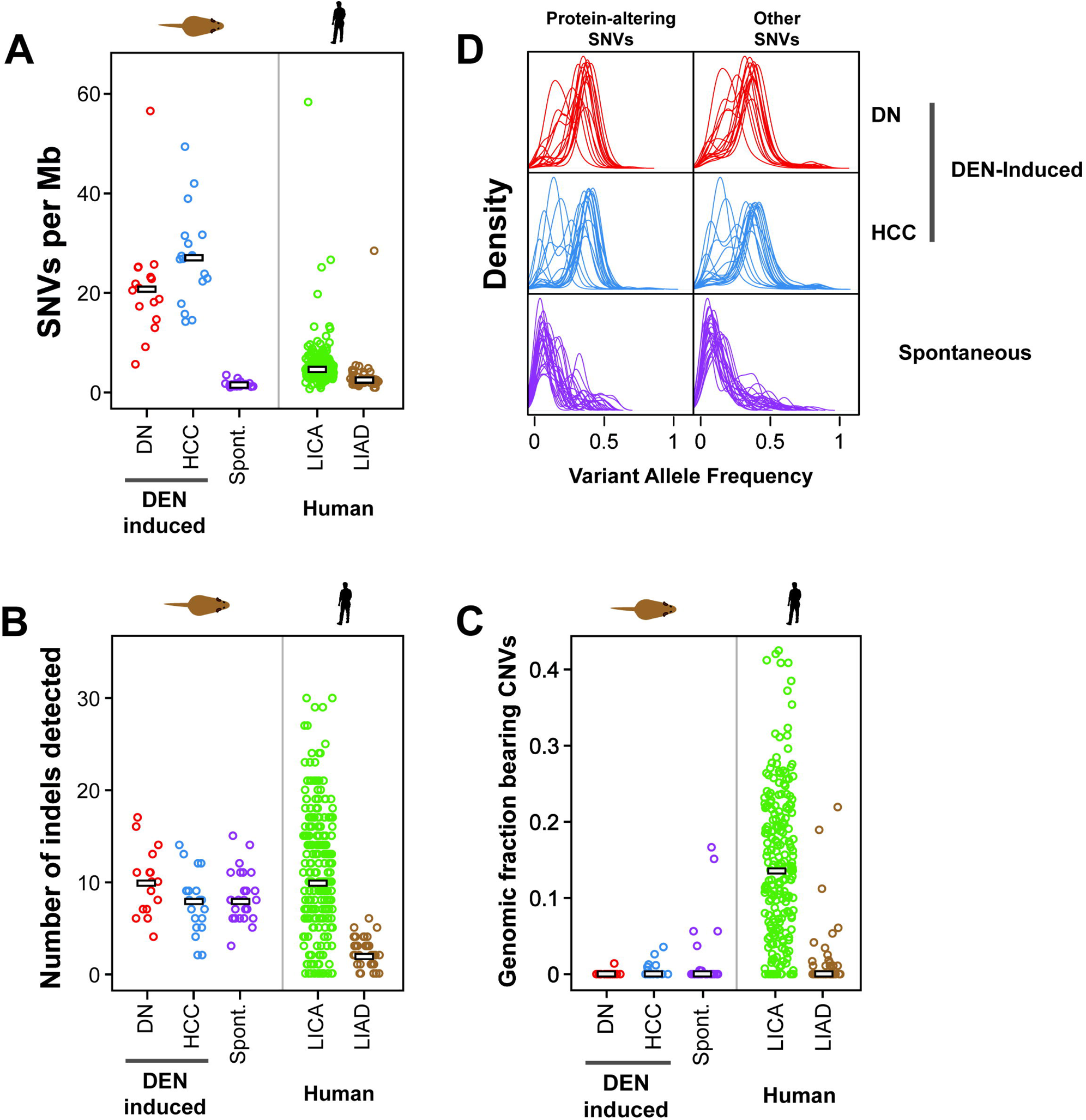
DEN-initiated neoplasms have a high SNV burden and few indels or copy number variations. (A) Estimated SNV mutation rates per megabase (Mb) in mouse and human liver tumour cohorts. The point mutation frequencies are shown for DEN-induced dysplastic nodules (DN, n=16) and hepatocellular carcinomas (HCC, n=18) and for spontaneous tumours (spont., n=25) arising in untreated mice. Previously reported human hepatocellular carcinoma (LICA, n=224) and hepatic adenoma (LIAD, n=38) samples are shown for comparison. Each point represents a single sample. Bars indicate median number of SNVs per Mb. (B) Comparison of frequencies of insertions and deletions (indels,1–50bp) in each cohort of mouse and human liver tumours. Each cohort had few indels, regardless of tumour histology or aetiology. Bars indicate median number of indels. (C) The fraction of the genome altered by cancer associated copy number variants (CNVs) in each cohort of mouse and human liver tumour samples. Mouse tumours had a lower fraction of their genomes present in CNVs larger than 10 Mb compared with human hepatocellular carcinomas (LICA). Bars indicate median genomic fraction with CNVs. (D) Distribution of variant allele frequencies (VAFs) in mouse liver tumours. Plots show the density of VAFs in DEN-induced dysplastic nodules (DN, n=16) and hepatocellular carcinomas (HCC, n=18) and in spontaneous tumours arising in untreated mice (n=25). Each line represents the distribution of VAFs from a single tumour. Separate plots are shown for SNVs classified as either protein-altering or as other (intergenic, intronic, or protein-coding synonymous). DEN-initiated tumours typically have higher VAFs (mean 0.32) than spontaneous neoplasms (mean 0.14).

The different aetiologies of the murine neoplasms may also explain their distinct SNV allele frequencies (Fig. 3D). The SNVs found in DEN-initiated tumours had a much higher variant allele frequency (VAF) than those found in spontaneous tumours (0.32 versus 0.14, on average, p-value 1.5 × 10^−5^). Spontaneous neoplasms carried many low abundance SNVs, and non-synonymous variants appear to be preferentially selected as a subset of these SNVs had increased VAFs (Fig. 3D). One likely explanation for this is the expansion of cells with acquired driver gene mutations (see also below). The uniformly high VAFs in carcinogen-initiated tumours is likely due to the single large burst of mutagenesis upon DEN exposure in the originating cell (Fig. 1B); the consequently high VAF might partially mask later acquisitions of driver mutations, selection and outgrowth of subclones.

### Distinct carcinogen imprint on the exome of DEN-induced neoplasms

All categories of somatic base substitutions were found in the exomes of DEN-initiated neoplasms, although C:G to G:C transversions were rarely detected (Fig. 4A; Table 1). Compared with the point mutations seen in untreated mice, DEN exposure resulted in an increase in transition and transversion events at A and T base pairs across the exome. These base substitutions are consistent with the persistence and mutagenicity of unrepaired alkylated thymidine lesions formed by metabolically activated DEN (11). T:A to A:T transversions and T:A to C:G transitions have been reported previously as predominant types of mutations induced by DEN, although these studies were limited to the sequencing of specific endogenous cancer genes or of surrogate genes in transgenic mouse mutation assays (12, 34).

**Table 1.**
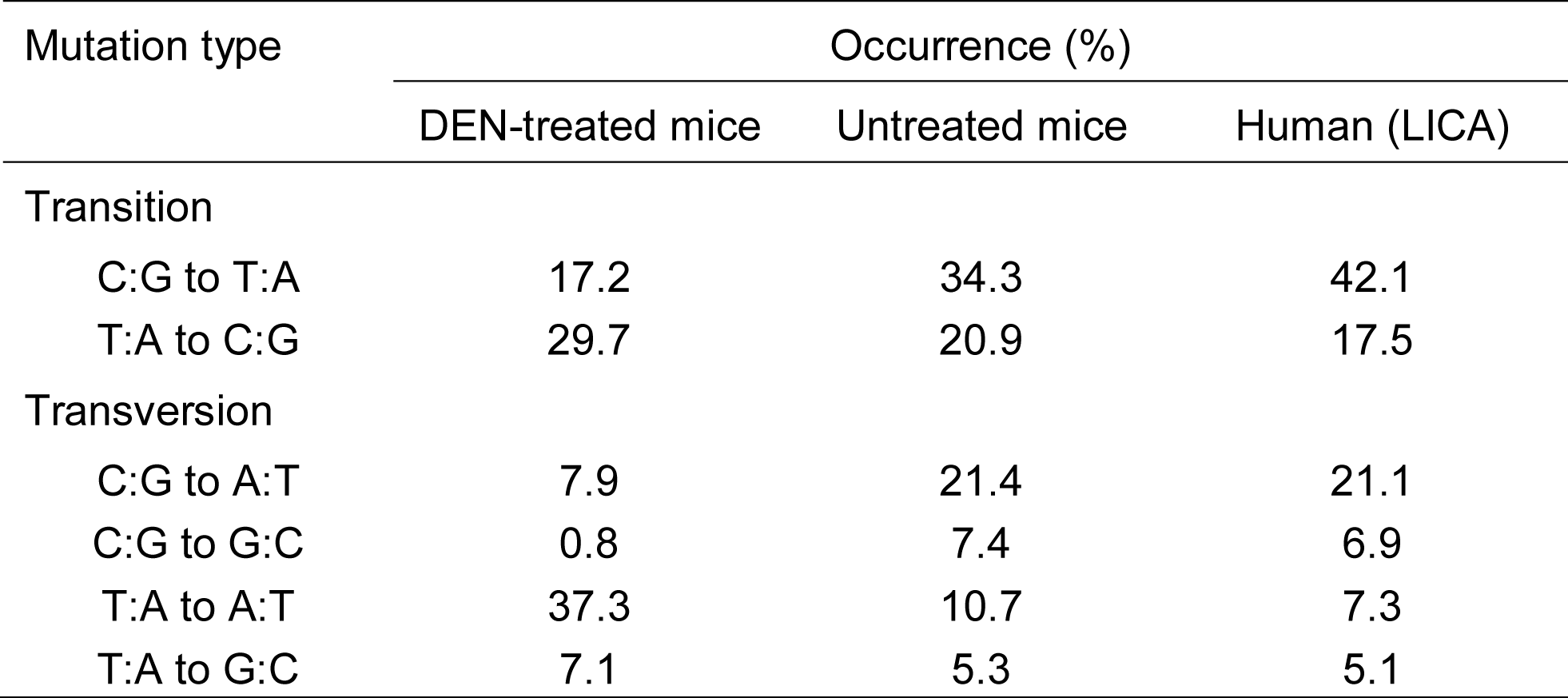
Mutation spectra in liver tumours from DEN-treated (n=34) and untreated (n=25) mice and in human HCC related to alcohol and metabolic syndrome (LICA) (n=224).

| Mutation type | Occurrence (%) |  |  |
| --- | --- | --- | --- |
|  | DEN-treated mice | Untreated mice | Human (LICA) |
| Transition |  |  |  |
| C:G to T:A | 17.2 | 34.3 | 42.1 |
| T:A to C:G | 29.7 | 20.9 | 17.5 |
| Transversion |  |  |  |
| C:G to A:T | 7.9 | 21.4 | 21.1 |
| C:G to G:C | 0.8 | 7.4 | 6.9 |
| T:A to A:T | 37.3 | 10.7 | 7.3 |
| T:A to G:C | 7.1 | 5.3 | 5.1 |

**Figure 4.**
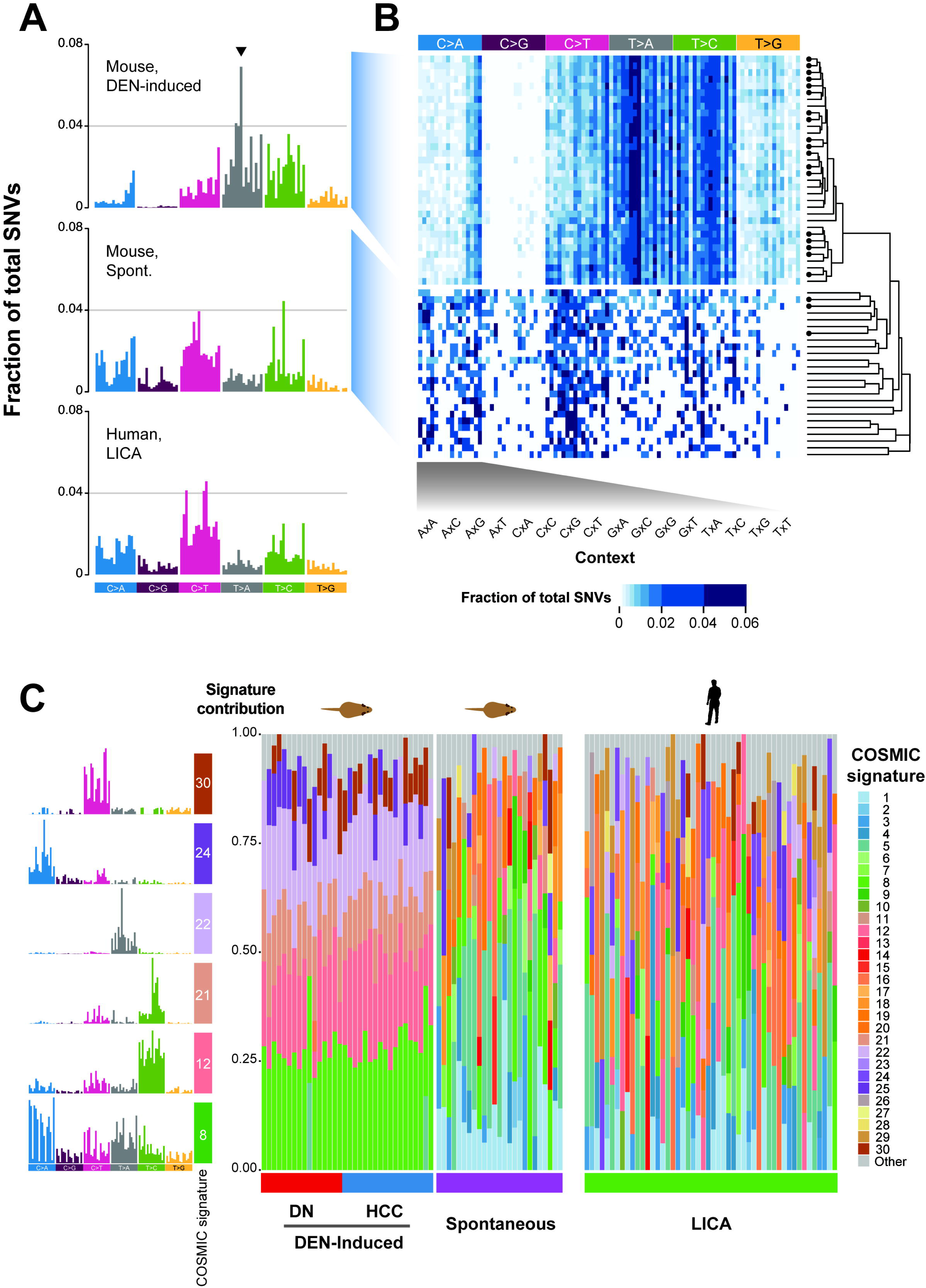
The exomes of DEN-initiated tumours have distinct and reproducible mutational profiles. (A) Frequencies of substitution mutations in mouse and human liver tumour cohorts. Mutational profiles are shown for DEN-induced mouse tumours (combined dysplastic nodule (DN) and hepatocellular carcinoma (HCC) samples, n=34); for spontaneous tumours (spont., n=25) arising in untreated mice; and for human liver tumours (LICA, n=50). The profiles are displayed using the 96 substitution classification, which is defined by reporting the specific base substitution combined with the immediate neighbouring 5’ and 3’ nucleobases. The arrow indicates one example of a trinucleotide context mutational bias observed in DEN-initiated tumours. (B) Heat map of the occurrence of mutational profiles of individual mouse tumours samples (rows) classified by substitution and trinucleotide context (columns). The right panel shows a phylogenetic tree quantifying the clustering observed from the individual mouse sample mutational profiles. A circle indicates a HCC sample; no circle indicates a DN sample. Neoplasms clustered by mutational profile, revealing a clear grouping into DEN-induced versus spontaneous tumours. (C) Mutational portraits of individual mouse and human liver tumours reconstructed using COSMIC mutational signatures. Each column shows the composition of signatures in an individual sample. DEN-induced mouse neoplasms showed reproducible portraits largely composed of six component COSMIC signatures (shown in the left panel). In contrast, mouse spontaneous and human tumour portraits were more heterogeneous.

The 5’ and 3’ nucleobases adjacent to point mutations in mouse DEN-induced tumours showed a complex pattern of biases. For example, we found a distinctive signature in DEN-initiated neoplasms where T:A to A:T transversions occurred more frequently when the T (or A) was preceded by a C (or A) and followed by a T (or G). Hierarchical clustering on the 96 possible trinucleotide substitution contexts showed a consistent mutational pattern shared among both dysplastic nodules and HCCs arising in C3H inbred mice exposed to DEN (Fig. 4B). The mutational patterns of tumours arising spontaneously in untreated C3H male mice clustered separately, highlighting the distinct mutational pattern of the carcinogen-initiated neoplasms (Fig. 4B).

Mathematical modelling of mutational processes in human cancer has defined over 30 mutational signatures, several of which are associated with exposure to specific environmental mutagens (35). We used these COSMIC signatures to computationally determine the composition of signatures which most accurately reconstructed the mutational profile of each mouse liver neoplasm (27). The resulting mutational portraits of the 34 DEN-initiated neoplasms were notably similar (Fig. 4C). The majority were largely composed of six reported COSMIC signatures: 8, 12, 21, 22, 24 and 30. Interestingly, signatures 12, 22 and 24 have been observed in human liver cancers, with signatures 22 and 24 reported to be associated with exposure to an exogenous mutagen, aristolochic acid or aflatoxin, respectively (6, 7, 9). The aetiologies of signatures 8, 12, 21 and 30 are currently unknown, although it has been speculated that the transcriptional strand bias reported in signatures 8 and 12 may reflect the involvement of transcription-coupled repair acting on bulky DNA adducts due to exogenous carcinogens (35). The six-signature mutational portrait that is characteristic of DEN-induced neoplasms is distinct from the mutational portraits of the 25 tumours arising spontaneously in untreated C3H male mice (Fig. 4C). The latter had a far more heterogeneous composition of individual mutational COSMIC signatures. We also carried out a similar analysis for human HCCs using the exome sequences of 50 randomly selected samples from the ICGC LICA-FR cancer genome project (6). The mutational portraits of these human liver cancers also had heterogeneous compositions of individual COSMIC mutational signatures (Fig. 4C). In sum, the mutational portraits of DEN-induced mouse tumours are remarkably homogeneous and reproducible, particularly in comparison with the diversity found within a typical cohort of human HCCs.

### Activating mutation of *Hras* is the most common driver of DEN-induced hepatocarcinogenesis in C3H mice

The neoplasms which arose following carcinogen exposure carried a high mutational load in their exomes. For example, each DEN-initiated dysplastic nodule had an average of 583 somatic SNVs in its protein coding sequence, compared with 26 SNVs in an average spontaneous neoplasm. As expected, 72% of these point mutations are predicted to be non-synonymous, with no detectable bias in the distribution of missense, nonsense and splice site mutations (Fig. 5A). Within our cohort of 34 carcinogen-initiated dysplastic nodules and HCCs, we have detected potential coding changes in 9222 genes (data not shown).

**Figure 5.**
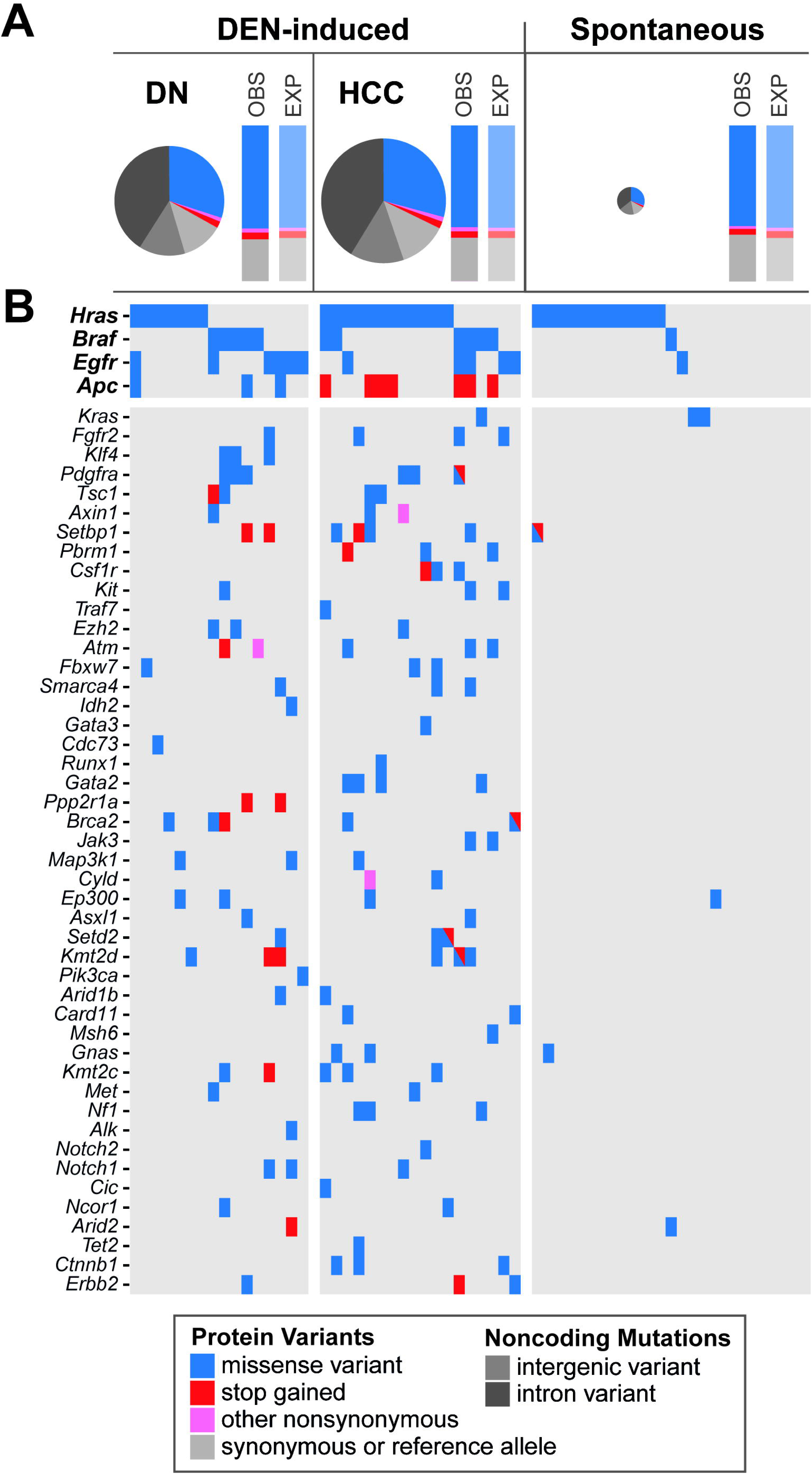
DEN-initiated and spontaneous mouse liver tumours carry recurrent activating mutations in *Hras*, but only carcinogen-induced tumours acquire a diversity of consequential SNVs in many cancer genes. (A) Proportions of predicted protein coding and non-coding variants in DEN-induced and spontaneous liver tumours. Pie charts show the proportions of each variant type in DEN-induced dysplastic nodules (DN, n=16) and hepatocellular carcinomas (HCC, n=18) and in spontaneous liver tumours (n=25). The observed (OBS) distribution of each substitution type was as expected (EXP), regardless of tumour histology or aetiology. The total area of the pie charts reflects the median SNV load within each sample set; spontaneous tumours had extremely low mutational loads. (B) Predicted consequential mutations in oncogenes and tumour suppressor genes for individual tumours. Each column in the table is a mouse tumour sample and each row is a cancer gene showing the occurrence of non-synonymous substitutions found in individual samples. Only genes mutated in at least two samples are shown (see Supplementary Table 1 for the complete list in each sample of somatic non-synonymous mutations in cancer genes).

We sought evidence for putative driver genes of hepatocarcinogenesis in our mouse model by searching for enrichment of non-synonymous mutations in validated oncogenes and tumour suppressor genes (23). We identified cancer genes which carried non-synonymous mutations more frequently than expected, with additional weight being given to genes which had recurrent hotspot SNVs (Fig. 5B; Supplementary Table 1). This approach revealed that *Hras* is the predominant, although not obligatory, oncogenic driver of hepatocellular carcinoma in juvenile male C3H mice which have been administered a single dose of DEN. Over half of the DEN-initiated tumour samples harboured a non-synonymous mutation in the *Hras* proto-oncogene, almost exclusively an activating hotspot mutation in codon 61 (Fig. 6A; Supplementary Table 2). The most common missense variant in codon 61 caused a glutamine to arginine substitution and was an A:T to G:C transition in the second base, which is consistent with the formation by DEN metabolites of one of the major promutagenic adducts, O4-ethyl-thymine. The incidence of *Hras* mutation increased from 44% in dysplastic nodules to 67% of HCC samples, suggesting that cells with oncogenic *Hras* had a selective advantage during DEN-initiated hepatocarcinogenesis. The neoplasms which arose in our untreated male C3H mice also had a high prevalence of non-synonymous mutations in *Hras* (48%), although the mutation spectrum was different. Almost half of the point mutations were identical G:C to T:A transversions in codon 117 (Fig. 6A; Supplementary Table 2), causing a lysine to asparagine substitution which is predicted to activate Ras (36).

**Figure 6.**
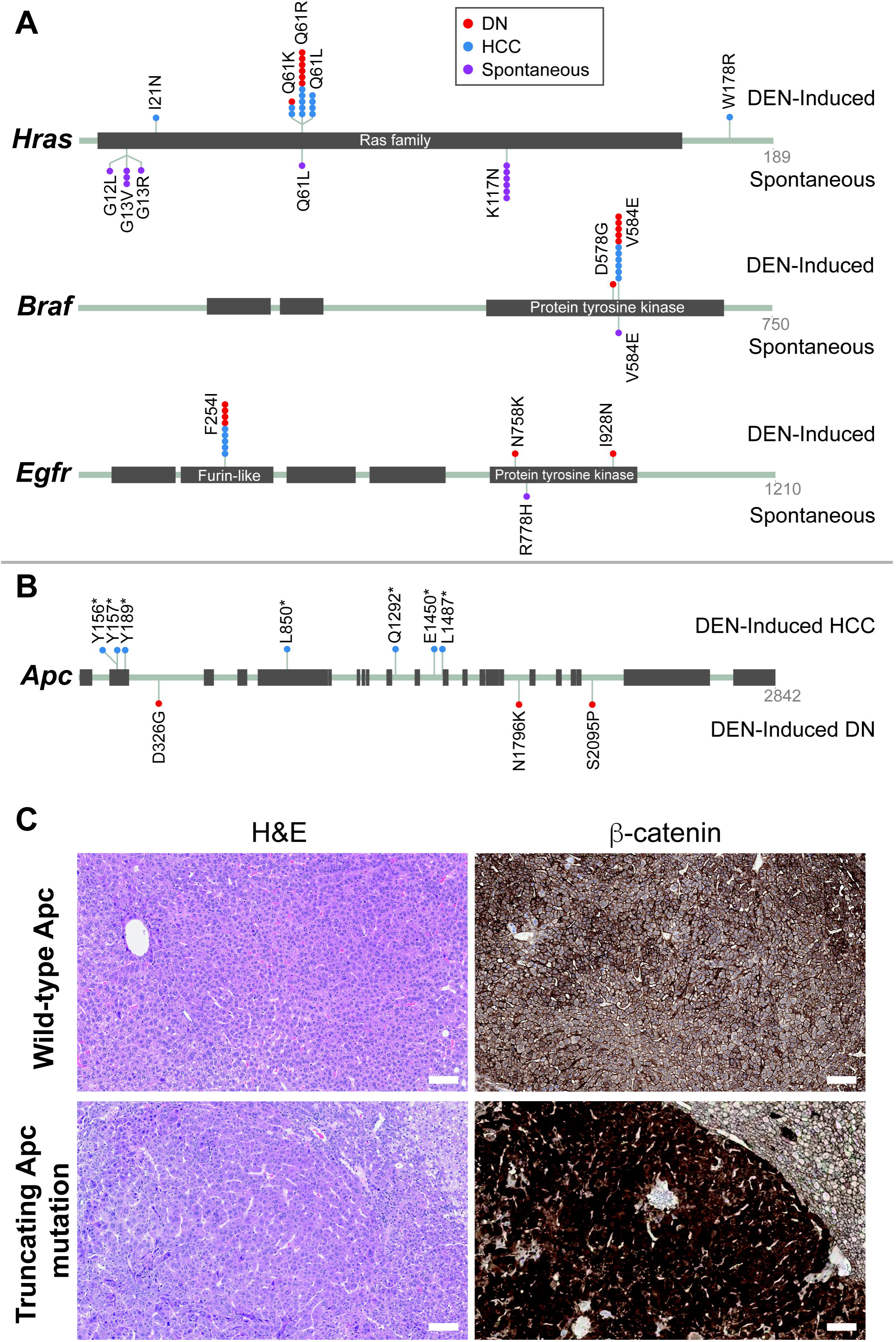
DEN-initiated and spontaneous mouse liver tumours acquire different recurrent mutations in putative driver genes. (A) Prevalence and location of somatic mutations in *Hras, Braf* and *Egfr* in DEN-initiated and spontaneous mouse liver tumours. Activating mutations in *Hras* were found at different hotspots in DEN-induced tumours (codon 61) compared with spontaneous tumours arising in untreated C3H mice (codon 117). Hotspot mutations in *Braf* (codon 5B4) and *Egfr* (codon 254) were recurrent in DEN-induced tumours, in contrast to spontaneous tumours which rarely carried non-synonymous SNVs in *Braf* or *Egfr*. (B) Prevalence and location of somatic mutations in *Apc* in DEN-induced hepatocellular carcinomas (HCC) compared with DEN-induced dysplastic nodules (DN). Truncating mutations in *Apc* were common in DEN-initiated tumours, exclusively in carcinoma samples. The spontaneous samples did not carry any non-synonymous mutations in *Apc* (not shown, see Supplementary Table 1). (C) Aberrantly elevated nuclear β-catenin protein expression in tumours with a nonsense mutation in *Apc*. Representative photomicrographs of serial tissue sections of DEN-induced HCCs. H&E staining demonstrates similar tumour morphology in tumours with wild-type *Apc* (upper panels) and in tumours with a nonsense *Apc* mutation (lower panels). IHC for β-catenin protein demonstrates aberrant strongly positive nuclear staining in *Apc* mutant HCC (lower panels). All scale bars = 100 μm. Original magnification x100.

Less frequently occurring oncogenic drivers of DEN-initiated hepatocarcinogenesis appear to be *Braf* and *Egfr* (Fig. 5B; Fig. 6A; Supplementary Table 2). Almost one third of tumours carried an identical activating hotspot mutation in *Braf:* an A:T to T:A transversion in codon 584, resulting in a valine to glutamic acid substitution in the kinase domain. We also identified a potentially activating hotspot missense mutation at codon 254 of the extracellular domain of *Egfr* in approximately one quarter of DEN-initiated tumours. An activating mutation in *Hras, Braf* or *Egfr* was present in every DEN-initiated neoplasm, although these mutations were very rarely found together in the same tumour. This apparent mutual exclusivity is likely because they can replace each other in terms of their oncogenic potential.

Every carcinogen-induced tumour carried non-synonymous SNVs in several *bona fide* cancer genes: on average we detected non-synonymous SNVs in five oncogenes and/or tumour suppressor genes in DEN-initiated tumours (range 1–11) (Supplementary Table 1). This considerable diversity of cancer genes that were mutated at low frequency after exposure to DEN limited our ability to detect any commonly mutated secondary drivers of DEN-initiated hepatocarcinogenesis. Nevertheless, we observed truncating mutations in *Apc* in 39% of HCC samples; nonsense mutations in *Apc* were not detected in the cohort of dysplastic nodules (Fig. 5B; Fig. 6B; Supplementary Table 2). The cancers bearing *Apc-* truncating mutations all showed aberrantly elevated levels of nuclear β-catenin (Fig. 6C), suggesting that loss of *Apc* function and disruption of the canonical WNT/β-catenin pathway can play a role in the progression to carcinoma in this model.

By comparison, spontaneous tumours from untreated mice contained few detectable point mutations in cancer genes (Fig. 5B; Supplementary Table 1). As previously discussed, approximately half were potentially driven by missense mutations activating the Ras signal transduction pathway. However, we could not unequivocally propose a driver gene for the remaining samples. The failure to detect other subtle mutations in potential driver genes may reflect a polyclonal composition of the spontaneous tumours and/or the involvement of other types of genetic or epigenetic alterations during tumorigenesis.

## Discussion

Chemically-induced mouse models of liver cancer are important tools widely used to study the molecular pathogenesis of human hepatocellular carcinoma (32). Over the last decade, large scale sequencing analyses of patient tumour samples have produced detailed profiles of the genetic aberrations found in human liver cancer genomes (4–9). It is important now to have similar descriptions of the genomic landscapes of the experimental mouse models used to inform the human disease (37, 38). Here we have described the mutational landscape of one of the most frequently used models of HCC, in which liver cancer is induced by a single injection of the genotoxin DEN into juvenile male mice.

Our strategy comparing spontaneously occurring liver tumours with those initiated by exposure to DEN allowed the direct comparison of the histopathology and genomic impact of carcinogen exposure. By controlling the initiating carcinogenic event the liver lesions in DEN-treated mice developed from early dysplastic nodules to carcinoma within a short, relatively consistent timeframe. The liver lesions arising in untreated mice arose at a much lower incidence and with a longer, more variable latency. Based on their histological appearance, liver tumours resulting from exposure to DEN were indistinguishable from those that arose spontaneously in untreated mice. This result parallels that found in human liver tumours, where heterogeneous molecular phenotypes can underlie HCC samples that are histologically similar. Indeed, these murine dysplastic lesions and carcinomas mimicked the histological features of their corresponding human tumours. In sharp contrast, however, this similarity was not seen in mutational landscapes: the exomes of DEN-induced neoplasms clearly reflected the DNA damage caused by chemical carcinogenesis.

DEN-induced tumours carried a notably high burden of somatic mutations, which allowed us to conclusively demonstrate the independent evolution of multiple macroscopic tumours within an individual liver. The mutational frequencies were much higher than those seen in most adult human solid tumours, including HCC. Perhaps not surprisingly, human lung and skin cancers that result from environmental exposure to potent mutagens are among the few human tumour types with mutational burdens similar to those we report here (23). Almost all of the DNA changes in the DEN-induced tumours were single base substitutions, consistent with the genotoxic action of DEN (11). Indeed, we confirmed that one of the major pro-mutagenic adducts caused by short-lived DEN metabolites was generated rapidly in centrilobular hepatocytes. It is likely that most of the genetic damage in tumours arising in livers exposed to DEN occurs when the originating hepatocytes are exposed to the carcinogen. In contrast to the elevated SNV levels, we did not find any evidence that DEN-induced cancer genomes have gross widespread alterations in chromosomal structure; we detected very few insertions, deletions or copy number variants in the exomes of DEN-induced tumours. This combination of a high exome-wide SNV burden with a paucity of copy number alterations has been observed in human cancers (39) as well as other carcinogen-induced mouse models of cancer (37).

Exposure to DEN left a common mutational imprint in the tumour exomes of treated mice. Indeed, the same small subset of reported signatures of mutational processes was readily identified computationally in every DEN-induced tumour. Notwithstanding this common imprint, each individual exome, including those of neoplasms arising within the same liver, carried a unique combination of somatic base substitutions. The majority of these SNVs are likely to be passenger mutations. However, we could identify four recurrently mutated genes that are putative oncogenic drivers of hepatocellular carcinoma in DEN-treated C3H male mice: *Hras, Braf, Egfr* and *Apc*.

The main genetic trait of DEN-initiated tumours is acquisition of mutations which deregulate signalling cascades involved in cell proliferation and survival. Over 80% of DEN-initiated tumour samples carried an activating hotspot driver mutation in either *Hras* or *Braf*. The remaining ~20% of samples carried a potentially activating hotspot mutation in *Egfr*, one of the upstream receptor tyrosine kinases that can regulate the Ras signalling pathway. This suggests that constitutive activation of the Ras/Raf/MEK/ERK signal transduction pathway is a hallmark feature in this mouse model of liver cancer.

Activation of the *Hras* proto-oncogene is frequently reported in both spontaneous and chemically-induced liver tumours in mice (12). However, the incidence and spectrum of *Hras* mutations appears to be strongly influenced both by the mouse strain used, as well as by the type and dose of chemical and experimental induction protocol employed. Indeed, even between spontaneous and treatment-induced tumours in C3H mice, we observed a difference in the location of the hotspot activating mutation in *Hras* (codon 117 versus codon 61, respectively). The mutational activation of *Braf* is also reported to be influenced by the mouse strain and appears to be related to strain susceptibility to hepatocarcinogenesis (13). As expected, we observed a lower frequency of mutations in *Braf* compared to *Hras* in the DEN-initiated liver tumours in the highly susceptible C3H mouse strain.

*Apc*, the third putative driver we identified, may be involved in progression to carcinoma in the DEN liver cancer model; we detected recurrent *Apc* truncating mutations only in HCC and never in dysplastic nodules. Activating β-catenin *(Ctnnb1)* mutations have been previously implicated in progression to carcinoma, but in a two stage model where DEN is given as the initiator followed by treatment with phenobarbital as a tumour promoter (14). In contrast with other reports using mice treated with DEN alone (that is, in the absence of a promoter) (40), we did observe (i) disruption of the canonical Wnt/β-catenin pathway, and (ii) that this disruption was caused primarily by loss-of-function mutations in *Apc* and consequent aberrant nuclear expression of β-catenin.

Aside from these four driver genes, there were no other *bona fide* cancer genes that were recurrently mutated. Instead, we saw a diversity of low incidence, non-synonymous point mutations in numerous oncogenes and tumour suppressors, consistent with the known mechanism of mutagenesis by DEN. Specifically, the introduction of a large mutagenic SNV burden stochastically across the genome resulted in heterogeneity at the level of driver gene mutations in the resulting tumours. However, it is also possible that common driver genes could have been dysregulated by alternative genetic or epigenetic processes during tumorigenesis.

Next generation sequencing of hundreds of human liver tumours has defined several cellular processes and pathways that are recurrently mutated in HCC (4–9). However, there appears to be a wide range of driver mutations that may underlie the dysregulation of these pathways during liver tumorigenesis. The DEN-initiated mouse model of HCC we described here replicates the Ras/MAPK signalling dysregulation found in a small subset of human HCC genomes, although activating mutations in the *RAS* family members have rarely been observed in human clinical samples. Perturbations in WNT/β-catenin signalling have been found in a large proportion of human HCC, and this is a common feature in progression to carcinoma in our DEN mouse model.

There are currently several mouse models of liver cancer, each of which recapitulates specific genetic, molecular, and/or histological features of the human disease (41). Here, we have described how chemical carcinogen-induced HCC, which is one of the more widely used mouse models of human HCC, introduces widespread single-base variations into the hepatocyte genome. Controlling both the starting inbred genome and the initiating carcinogenic event generates matched, independently-evolved tumour samples within a consistent time frame. The exomes of these liver tumours carry a large mutational burden of single nucleotide variations that reflect the imprint left after exposure of the originating hepatocyte to the carcinogen. The tumours generated by diethylnitrosamine in this mouse model are driven by activating mutations in *Hras, Braf* or *Egfr*, with a role for deregulation of Wnt/β-catenin signalling in the progression to carcinoma.

Our study demonstrates how the application of exome sequencing on carefully designed cohorts can reveal novel insights into widely used mouse models of liver cancer. Such oncogenomic descriptions will deepen our understanding of the advantages and limitations of preclinical *in vivo* models and thereby inform the selection of the most appropriate models to study human liver cancer.

## Acknowledgements

We thank the following CRUK Cambridge Institute core facilities for their vital contributions: Biological Resources, Pre-clinical Genome Editing, Histopathology & ISH, Research Instrumentation and Genomics. In particular, we acknowledge technical support from Lisa Young, Steven Kupczak, Maureen Cronshaw, Paul Mackin, Yi Cheng, Lena Hughes-Hallett, Angela Mowbray, Jodi Miller and Leigh-Anne McDuffs, and advice from James Hadfield, Matthew Eldridge, Ruben Drews and Oscar Rueda. We also thank Susan E Davies for assistance with histopathology.

## Supplementary Information

### Materials and Methods

#### Mouse models of hepatocarcinogenesis

All animal experimentation was carried out in accordance with the Animals (Scientific Procedures) Act 1986 (United Kingdom) and with the approval of the Cancer Research UK Cambridge Institute Animal Welfare and Ethical Review Body. C3H/HeOuJ strain mice were obtained from Charles River Laboratories and maintained using standard husbandry. Mice were housed in Techniplast GM500 Mouse IVC Green Line cages in a room with 12 hour light / 12 hour dark cycle. When possible, animals were group housed. Cages contained aspen bedding and the following cage enrichments: nesting material, aspen chew stick and cardboard tunnel. Mice had *ad libitum* access to water and food (LabDiet 5058).

Liver tumours were chemically induced in male mice aged 14–16 days by a single intraperitoneal injection of DEN (Sigma-Aldrich N0258; 20mg/kg body weight) diluted in 0.85% saline. To assess the immediate response to DEN, some mice were humanely killed at 4, 8, 12, and 24 hours after DEN injection. Liver tumour samples were collected from DEN-treated mice aged for 24 to 40 weeks after injection. A separate cohort of untreated male mice was aged for 37 to 76 weeks to develop spontaneous liver neoplasms.

#### Tissue collection, processing, and staining

Ear and tail samples from untreated mice were flash frozen for use as a normal genome reference. Liver tumours were macroscopically identified and isolated. Small nodules were flash frozen in liquid nitrogen for subsequent DNA extraction. Nodules of sufficient size (>2mm diameter) were bisected; one half was flash frozen in liquid nitrogen for DNA extraction and the other half was fixed in neutral buffered formalin for 24 hours, transferred to 70% ethanol, machine processed (Leica ASP300 Tissue Processor) and paraffin embedded. All formalin-fixed paraffin-embedded sections were 3 μm in thickness.

Formalin-fixed paraffin-embedded tissue sections were stained with haematoxylin and eosin for morphological assessment and using the Gomori’s method for reticular fibres to assess liver architecture. Histochemical staining was performed using the automated Leica ST5020; mounting was performed on the Leica CV5030. Immunohistochemistry was performed on formalin-fixed paraffin-embedded sections using the Bond Polymer Refine Kit (DS9800, Leica Microsystems) with DAB enhancer (Leica Biosystems, AR9432) on the automated Bond platform. Immunohistochemistry was performed using antibodies against β-catenin (BD Biosciences, 610154, 1:100 dilution); phospho-histone H2AX (Merck Millipore, MABE205, 1:5000); O^6^-ethyl-2-deoxyguanosine (ER6, Squarix Biotechnology, SQM001.1, 1:200); and Ki67 (Bethyl Laboratories, IHC-00375, 1:1000). Heat-induced epitope retrieval was performed for 20 minutes at 100°C on the Bond platform with sodium citrate. For the anti-O^6^-ethyl-2-deoxyguanosine staining the peroxidase block was done post primary antibody incubation and the post primary component of the polymer kit was substituted with a rabbit anti-rat secondary antibody (Bethyl Laboratories, A110–322A, 1:250). All tissue sections were scanned using Aperio XT (Leica Biosystems); H&E-stained tissue section images are available at BioStudies archive at EMBL-EBI under accession S-BSMS4.

Quantification of nuclear staining for O^6^-ethyl-2-deoxyguanosine and phospho-histone H2AX was done using ImageScope software (Leica Biosystems). On average 505,000 cells from one cross-section per sample were evaluated (range: 290,000 - 690,000). Percent of positive-staining nuclei were plotted using the ggplot2 package in RStudio and statistical significance was calculated using the Welch two sample t-test.

#### Histological classification and selection of liver tumours

Tissue sections were blinded and assessed twice by a histopathologist; discordant results were reviewed by an independent hepatobiliary histopathologist. Tumours were classified according to the International Harmonization of Nomenclature and Diagnostic Criteria for Lesions in Rats and Mice (INHAND) guidelines (17). Dysplastic nodules have an expansile growth pattern causing compression of adjacent hepatic parenchyma, loss of normal lobular architecture (irregular reticulin fibre staining), nuclear atypia, and may show increased proliferation (increased Ki67 staining). Hepatocellular carcinomas are characterised by thickened trabeculae (loss of reticulin fibre staining), pseudoglandular structures, more marked cellular atypia, increased nuclear to cytoplasmic ratios, higher proliferative index (markedly increased Ki67 staining) and an infiltrative growth pattern.

Tumours with sufficient tissue for histological classification were selected for whole exome sequencing experiments if they met the following criteria: (i) diagnosis of either DN or HCC, (ii) homogenous tumour morphology, (iii) tumour cell percentage >80%, and (iv) adequate tissue for DNA extraction. Neoplasms with extensive necrosis, mixed tumour types, a nodule-in-nodule appearance (indicative of an HCC which had arisen within a DN), or contamination by normal liver tissue were excluded.

#### DNA isolation and whole exome sequencing

Genomic DNA from liver tumours was isolated using the AllPrep DNA/RNA mini kit (Qiagen, 80204). Genomic DNA from ear/tail samples was extracted with the DNeasy blood & tissue kit (Qiagen, 69506). Exome capture libraries were prepared following the instructions of the SureSelectXT mouse all exon target enrichment system (Agilent Technologies, 5190–4641). Briefly, 3μg DNA were sheared to 150–300bp fragments using the Covaris S220 system. Libraries were generated with the SureSelectXT reagent kit (G9611A), hybridised and enriched according to the manufacturer’s instructions. During this process all clean-up steps were performed with Agencourt AMPure XP beads (Beckman Coulter, A63880), all DNA concentration measurements were done with the Qubit fluorometric method (ThermoFisher) and library quality controls were run on the Agilent Bioanalyzer 2100 (DNA1000, DNA high-sensitivity). Finally, the exome libraries were quantified by real-time PCR using the Kapa library quantification kit (KapaBiosystems) on the QuantStudio 6 Flex (Applied Biosystems) before pooling and sequencing 125bp paired-end reads on an Illumina HiSeq2500.

#### Sequencing read alignment

Sequencing reads were aligned to the C3H_HeJ_v1 mouse genome assembly (Ensembl release 90 (18)) with bwa (versions 0.6.1 or 0.7.12 (19)). Aligned reads were annotated to read groups using the picard tool AddOrReplaceReadGroups and minor annotation inconsistencies corrected using the picard CleanSam and FixMateInformation tools (picard version 1.124; http://broadinstitute.github.io/picard). The bam files for each sample were merged together and duplicates reads were then identified using the picard MarkDuplicates tool. Sequencing coverage was assessed using samtools (version 1.1 (20)). 95% of genome regions predicted to represent coding sequence were found to be covered to a read depth of 20 or greater. Raw sequencing reads are available from European Nucleotide Archive at EMBL-EBI under accession PRJEB19083. Aligned reads for human samples from the LICA-FR and LIAD-FR cohorts were downloaded from the European Genome-phenome Archive at EMBL-EBI (accessions EGAD00001000131, EGAD00001001096 and EGAD00001000737).

#### Variant calling

A pooled normal sample set was generated by combining all control sample reads and then subsampling these reads to match the mean coverage achieved for the control and tumour samples. Single nucleotide and indel variants were called using Strelka (version 1.0.14 (21)) using the recommended configuration for bwa-aligned reads and setting the isSkipDepthFilters flag for improved calling on exome-seq data. Variant calls were combined into a merged set using bcftools (version 1.1 (20)). Predicted coding sequence changes due to the SNVs were annotated by comparing the polypeptide sequences coded by the reference and variant alleles.

SNVs were subjected to multiple filtering steps. Firstly, low-confidence SNV calls were identified and removed by applying the recommended filters for Strelka output from the gatk-tools package (version 0.2 (42); https://github.com/crukci-bioinformatics/gatk-tools). In particular, variants were filtered based on low mapping and base quality scores, proximity to alignment ends, and low absolute read counts. A VCF file is available from European Nucleotide Archive at EMBL-EBI under accession ERZ389577. Secondly, SNVs with an allele frequency of less than 2.5% were omitted to eliminate possible cross-contamination due to observed low levels of sequence read index mis-assignment during Illumina sequencing (James Hadfield, personal communication).

SNV call rates were estimated by fitting the number of variants detected at a range of sequencing read depths and extrapolating to determine the expected call rate at saturated sequencing coverage. Variant allele frequencies were calculated from read counts for reference and variant alleles, excluding those reads having a MAPQ score less than 5. Confidence intervals for allele frequency estimation were estimated by applying a normal distribution approximation to bootstrap-resampled frequency estimates (R boot package version 1.3 (43)).

Autosomal copy number variations (CNVs) were called with CNVkit (version 0.7.2 (22)) using default parameters. CNV regions were filtered to remove low-confidence regions where the null hypothesis (i.e., unchanged copy number) fell within the 95% confidence interval. Further filters removed CNVs where the absolute log fold change in copy number was smaller than 0.25. CNV regions located within 10kb of each other were merged, and the resulting regions finally filtered to remove regions smaller than 10Mb.

#### SNV validation

Non-synonymous SNVs in the cancer driver genes of interest, *Hras, Braf, Egfr* and *Apc*, were validated using conventional Sanger sequencing. The majority (87%) were validated using this method. The remaining SNVs could not be validated by Sanger sequencing for technical reasons, for example, there was no sample available or because the variant allele frequency was too low to allow confident detection by this method. These SNVs were checked by visual inspection of the aligned reads and were called validated if the total variant reads were greater than ten. The method used for each SNV validation is noted in Supplementary Table 3.

#### Independence of tumour evolution

For phylogenetic and mutational signature analyses (see below) the SNV list was filtered to remove those genome regions which were not covered by at least 20 non-duplicated reads (i.e. coverage of 20x or above) across all samples. This yielded a data set in which samples may be reasonably assumed to bear homozygous reference alleles at the vast majority of loci for which Strelka does not call a variant. Protein coding, exonic, and genic regions represented 42.6%, 53.7% and 92.0% of these 20x covered genomic regions, respectively.

A phylogenetic tree describing the development of the tumours within our cohorts was constructed in R using the ape package (version 3.5 (24)). Pairwise distances between samples were calculated as the number of genomic loci at which the sample genotypes differ. Trees were constructed using a neighbour-joining algorithm (25).

#### Mutational signature analysis

Analysis of mutational signatures was constrained to just those regions covered to at least 20x in all samples. The distributions of 5' and 3' nucleotides flanking the called SNVs were calculated directly from the reference genome. Direct comparison between human and mouse signatures was facilitated by normalising C3H/HeJ nucleotide context distributions using the ratios of known trinucleotide prevalences in C3H/HeJ and human genomes, as calculated for the 20x covered regions for each genome.

The proportions of COSMIC mutational signatures (26) represented in the mutational profile from each sample was calculated using the R package deconstructSigs (version 1.8.0 (27)).

#### Identification of significantly mutated cancer-related genes

Variants were annotated as to their likely effect on coding sequence by comparing their predicted polypeptide sequences to those from the Ensembl release 90 C3H_HeJ_v1 reference. SNV calls which were shared between tumours taken from the same mouse (i.e., SNVs which could be simply ascribed to germline variation) were filtered prior to analysis of mutated genes.

Cancer-related genes bearing above expected levels of non-synonymous mutations (both across the entire gene and recurring at specific loci) were identified using the following procedure. The gene list to be analysed was constrained to the listing of oncogenes and tumour suppressor genes described by Vogelstein *et al*., 2013 (23). The total mutation load (nonsense, missense, frame shift, splice site) within coding and splice-site regions was used to calculate the probability that mutations had occurred purely by stochastic mutational processes. Within each gene, the count of mutations at each variant locus was fitted to a Poisson distribution assuming a background mutation rate calculated across all sequenced regions. The individual variant loci were combined at the gene level using a multinomial model using the R XNomial package (version 1.0.4; https://CRAN.R-project.org/package=XNomial). This model yielded log likelihood ratios from the observed and expected distributions, from which a gene-wise P-value was readily calculated. P-values were corrected for multiple testing across all genes using the Bonferroni method. The analysis was repeated imposing a gene filter derived from the Cosmic Cancer Gene Census (http://cancer.sanger.ac.uk/census/). This identified the same cancer-associated genes that were significantly mutated in the mouse neoplasms.

**Supplementary Table 1.** Somatic non-synonymous SNVs detected in cancer driver genes in DEN-induced and spontaneous mouse liver tumours. Individual liver tumour samples are arranged in columns using the mouse and tumour identification codes (DN: DEN-induced dysplastic nodule; HCC: DEN-induced hepatocellular carcinoma; spontaneous: tumour from an untreated mouse). Details of the SNVs in oncogenes and tumour suppressor genes are shown in the rows. SNVs are coded as “0/1” to indicate a detected heterozygous variant genotype, and “.” to indicate no variant call. Predictions of the likely consequences of mutations were incorporated as follows: genomic locations of the SNVs were lifted over from the C3H_HeJ_v1 genome assembly to the GRCm38 mouse assembly; the resulting loci were used to query PROVEAN, SIFT and COSMIC database tools.

**Supplementary Table 2.** Occurrence of non-synonymous SNVs in *Hras, Braf, Egfr* and *Apc* in dysplastic nodules (DN) and hepatocellular carcinomas (HCC) from livers of DEN-treated and untreated C3H mice.

| Gene | Mutation | Number of neoplasms with the specified mutation |  |  |  |
| --- | --- | --- | --- | --- | --- |
|  |  | DEN-initiated | DEN-initiated | Spontaneous | Spontaneous |
|  |  | DN<br>(n=16) | HCC<br>(n=18) | DN<br>(n=22) | HCC<br>(n=3) |
| *Hras* | G12L | 0 | 0 | 1 | 0 |
| *Hras* | G13R | 0 | 0 | 0 | 1 |
| *Hras* | G13V | 0 | 0 | 3 | 0 |
| *Hras* | I21N | 0 | 1 | 0 | 0 |
| *Hras* | Q61K | 1 | 2 | 0 | 0 |
| *Hras* | Q61L | 0 | 4<sup>a,c,d</sup> | 1 | 0 |
| *Hras* | Q61R | 6<sup>g</sup> | 5 | 0 | 0 |
| *Hras* | K117N | 0 | 0 | 6 | 0 |
| *Hras* | W178R | 0 | 1<sup>a</sup> | 0 | 0 |
| *Braf* | D578G | 1<sup>b</sup> | 0 | 0 | 0 |
| *Braf* | V584E | 5<sup>b</sup> | 6<sup>c,e,f</sup> | 1 | 0 |
| *Egfr* | F254I | 4 | 5<sup>d,e</sup> | 0 | 0 |
| *Egfr* | N758K | 1<sup>f</sup> | 0 | 0 | 0 |
| *Egfr* | R778H | 0 | 0 | 1 | 0 |
| *Egfr* | I928N | 1<sup>g</sup> | 0 | 0 | 0 |
| *Apc* | Y156* | 0 | 1 | 0 | 0 |
| *Apc* | Y157* | 0 | 1 | 0 | 0 |
| *Apc* | Y189* | 0 | 1 | 0 | 0 |
| *Apc* | spl?<sup>h</sup> | 0 | 1 | 0 | 0 |
| *Apc* | D326G | 1 | 0 | 0 | 0 |
| *Apc* | L850* | 0 | 1 | 0 | 0 |
| *Apc* | Q1292* | 0 | 1 | 0 | 0 |
| *Apc* | E1450* | 0 | 1 | 0 | 0 |
| *Apc* | L1487* | 0 | 1 | 0 | 0 |
| *Apc* | N1796K | 1 | 0 | 0 | 0 |
| *Apc* | S2095P | 1 | 0 | 0 | 0 |

<sup>a</sup> *Hras* W178R co-occurred with *Hras* Q61L in one DEN-initiated HCC sample.
<sup>b</sup> *Braf* D578G co-occurred with *Braf* V584E in one DEN-initiated DN sample.
<sup>c</sup> *Braf* V584E co-occurred with *Hras* Q61L in one DEN-initiated HCC sample.
<sup>d</sup> *Egfr* F254I co-occurred with *Hras* Q61L in one DEN-initiated HCC sample.
<sup>e</sup> *Egfr* F254I co-occurred with *Braf* V584E in two DEN-initiated HCC samples.
<sup>f</sup> *Egfr* N758K co-occurred with *Braf* V584E in one DEN-initiated DN sample.
<sup>g</sup> *Egfr* I928N co-occurred with *Hras* Q61R in one DEN-initiated DN sample.
<sup>h</sup> spl?, splice acceptor variant that might affect splicing.

**Supplementary Table 3.**
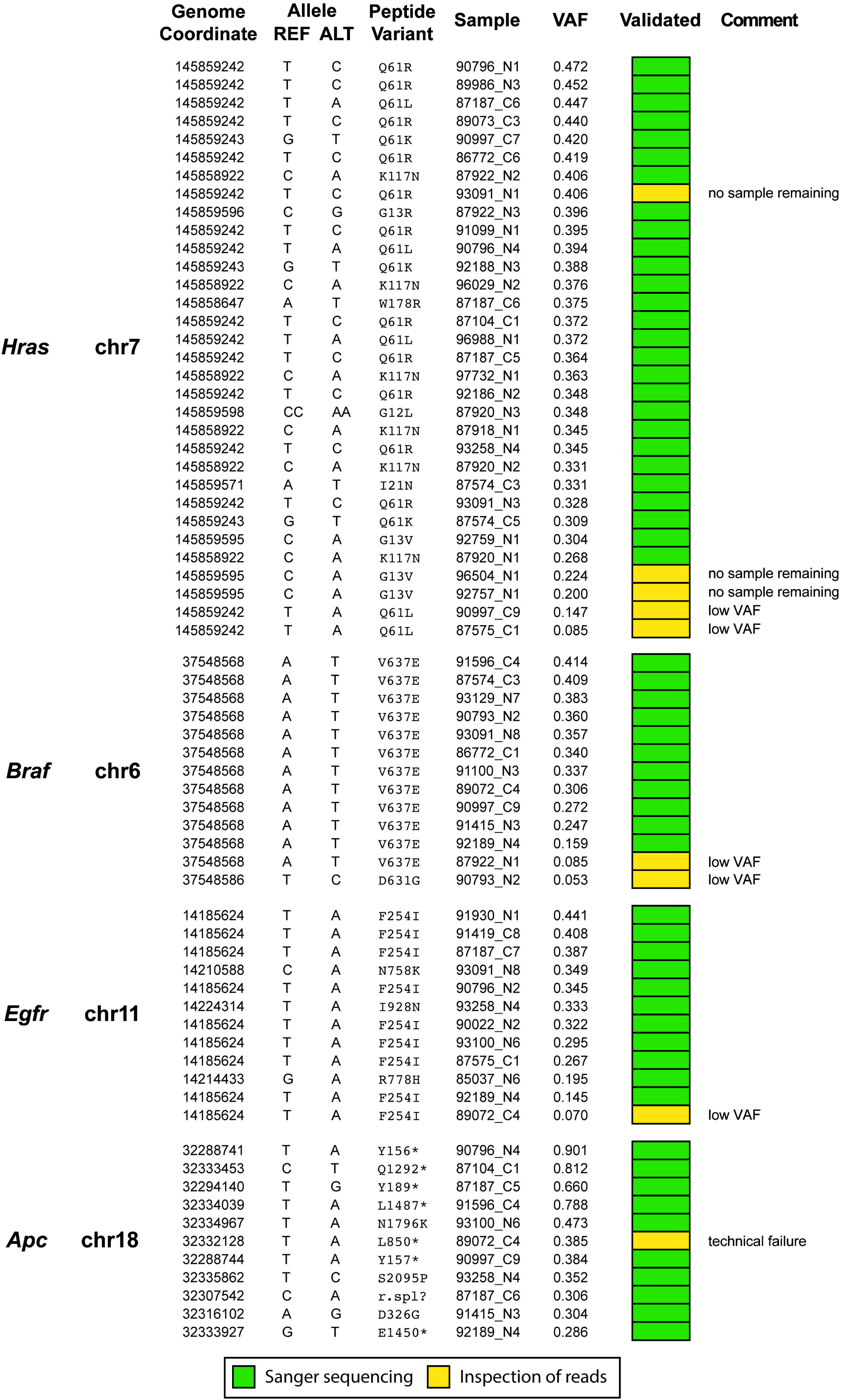
Non-synonymous SNVs in *Hras, Braf, Egfr* and *Apc* in mouse liver neoplasms. The tumour samples carrying a non-synonymous SNV in *Hras, Braf, Egfr* and/or *Apc* are listed. Details of the individual SNVs, including genome coordinate, nucleotide and peptide variant, variant allele frequency (VAF) and validation method, are shown for each liver tumour sample. Variants were validated by Sanger sequencing whenever possible, or by visual inspection of aligned reads

Author names in bold designate shared co-first authorship.

